# Intracellular carbon storage enables starvation survival in marine bacteria

**DOI:** 10.64898/2026.03.25.714165

**Authors:** Olesia Shlakhter, Yonathan Talmor, Sergey Malitsky, Leilah Otikovs, Amir Szitenberg, Einat Segev

**Affiliations:** Department of Plant and Environmental Sciences, Weizmann Institute of Science; Rehovot, 7610001, Israel; Department of Life Sciences Core Facilities, Weizmann Institute of Science; Rehovot, 7610001, Israel; Department of Chemical Research Support, Weizmann Institute of Science; Rehovot, 7610001, Israel; Ilana And Pascal Mantoux Institute for Bioinformatics, Weizmann Institute of Science; Rehovot, 7610001, Israel

## Abstract

Heterotrophic marine bacteria frequently experience fluctuations in carbon availability driven by phytoplankton dynamics. As a result, bacteria undergo repeated cycles of rapid growth during brief resource pulses followed by prolonged starvation. Yet the mechanisms that support bacterial survival during nutrient limitation remain poorly understood. Here, we investigate starvation survival in the algal-associated bacterium *Phaeobacter inhibens*. We show that cells remain viable for extended periods under carbon depletion while undergoing physiological and morphological changes. Using electron microscopy, metabolomics, and genetic approaches, we identify intracellular polyhydroxybutyrate (PHB) granules as a key factor supporting survival during starvation. PHB accumulates during growth and is progressively consumed under carbon limitation. Deletion of the PHB synthase gene (*phaC*) eliminates granule formation and reduces long-term viability. Comparative analyses show that the genetic capacity for PHB biosynthesis is widespread among members of the Roseobacter group, suggesting a conserved strategy among algal-associated bacteria. However, species lacking PHB also survive starvation, indicating that additional mechanisms contribute to persistence under nutrient limitation. Together, our results identify intracellular carbon storage as a central mechanism linking bacterial physiology to survival in fluctuating marine environments, and highlight the diversity of strategies shaping microbial community dynamics and carbon cycling in the ocean.

## Introduction

Heterotrophic bacteria in the ocean often live in close association with microalgae (Buchan et al. 2014). By attaching to algal cells or residing in their immediate surroundings, these bacteria can use organic compounds released during photosynthesis (Buchan et al. 2014; Thornton 2014). As a result, bacterial growth frequently tracks algal abundance and productivity. Because algae rely on light to drive carbon fixation, their metabolism follows daily light-dark cycles (Granum et al. 2002; Doblin et al. 2011; Leboulanger et al. 1997). Beyond daily oscillations, phytoplankton populations also fluctuate on seasonal scales. Periods of intense primary production and plentiful algal exudates alternate with phases of low photosynthetic activity and reduced carbon supply (Gasol et al. 2016). The diel and seasonal cycles of phytoplankton create a dynamic resource landscape for heterotrophic bacteria. When algal-derived substrates are abundant, copiotrophic bacteria consume the algal exudates and grow (Lauro et al. 2009; Song et al. 2017). However, when algal production declines, resources become scarce, and bacteria face starvation. Therefore, algal-associated bacteria must cope with recurring transitions between rapid growth and nutrient deprivation.

Large regions of the ocean are oligotrophic, characterized by low concentrations of nutrients and highly dilute pools of dissolved organic carbon (Morita 1988; Azam and Malfatti 2007). In such environments, heterotrophic bacteria rarely experience sustained resource abundance. Instead, they encounter episodic pulses of organic substrates that are rapidly consumed, followed by extended periods of scarcity. This feast-famine dynamic means that starvation is not an exceptional condition but rather a common physiological state for many marine bacteria (Moriarty and Bell 1993). Consequently, the ability to endure prolonged nutrient limitation is a key determinant of microbial survival and ecological success in marine ecosystems. This challenge is particularly relevant for bacteria that do not form specialized resistant structures such as spores or cysts, in which strategies for long-term survival remain only partly characterized (Novitsky and Morita 1978; Jones and Rhodes-Roberts 1981; Gray et al. 2019). Understanding how these bacteria persist between transient resource pulses is therefore central to explaining the structure and function of ocean microbial communities.

Bacteria use a range of molecular strategies that allow them to survive nutrient deprivation, but the underlying mechanisms were elucidated in a handful of model bacteria. In *Escherichia coli*, cells enter the stationary phase through major transcriptional and metabolic changes that reduce growth, increase stress resistance, shift central metabolism, and alter membrane composition (Kolter et al. 1993; Finkel 2006; Nyström 2004; S. Schink et al. 2022). Cells also exhibit morphological changes such as cytoplasm shrinkage (Shi et al. 2021). Studies in *Vibrio* and *Pseudomonas* species similarly report reduced cell size (Kjelleberg et al. 1983; Amy and Morita 1983) and a transition into low-activity, dormancy-like states (Mårdén et al. 1985; Morita 1982). Some bacteria store intracellular compounds such as glycogen (L. Wang and Wise 2011) or polyhydroxyalkanoates (PHAs) (Jendrossek and Pfeiffer 2014), which cells can later use when external nutrients become limiting. Consistent with this strategy, several marine bacteria, including *Sphaerotilus discophorus, Pseudomonas sp*., *Vibrio spp*. and *Dinoroseobacter shibae*, contain intracellular PHA granules, that have been suggested to support survival during starvation (Stokes and Parson 1968; Jones and Rhodes-Roberts 1981; Mårdén et al. 1985; H. Wang et al. 2014; Chien et al. 2007). Although starvation is common in the ocean, the molecular mechanisms that allow marine bacteria, particularly those associated with phytoplankton, to survive nutrient limitation remain poorly understood.

Here, we examine starvation survival in *Phaeobacter inhibens*, a widespread marine bacterium that forms natural associations with diverse microalgae (Segev et al. 2016; Majzoub et al. 2019). Using electron microscopy, targeted metabolomics and genetic approaches, we uncover a central role for intracellular carbon storage granules in survival during carbon starvation. We identify the biosynthetic pathway responsible for granule formation and demonstrate that it is broadly distributed among members of the Roseobacter group, indicating a conserved strategy among many algal-associated bacteria. At the same time, our findings suggest that bacteria outside the Roseobacter lineage rely on alternative mechanisms to endure starvation. These strategies remain largely unexplored and point to a broader diversity of survival solutions in marine microbial communities.

## Results

### *P. inhibens* bacteria survive carbon limitation during long-term cultivation

We first examined how *P. inhibens* bacteria respond to prolonged starvation. Cultures were grown in CNS medium supplemented with glucose as a carbon source at 18°C to reflect environmental conditions (see Materials and Methods) and were monitored over 30 days of incubation. We assessed growth using both optical density measurements at 600 nm (OD_600_) and colony-forming unit (CFU) counts. In the context of starvation, each method has inherent limitations: OD_600_ is influenced by cell size and optical properties (Mira et al. 2022; Myers et al. 2013) and cannot distinguish between live and dead cells, whereas CFU counts provide direct evidence of viability but can be affected by multicellular structures (Segev et al. 2015). Therefore, using both methods provides a comprehensive estimate of cell abundance.

Our data show that after one week of incubation, cultures entered the stationary phase and no longer exhibited increasing cell counts (Fig. 1). The bacterial population remained viable over 30 days, with stable counts above 10^8^ CFU/ml. Based on the observed growth dynamics, we defined specific time points to represent distinct growth phases: day 3, exponential phase; day 7, early stationary phase; day 10, early starvation; and day 17, late starvation.

**Figure 1.**
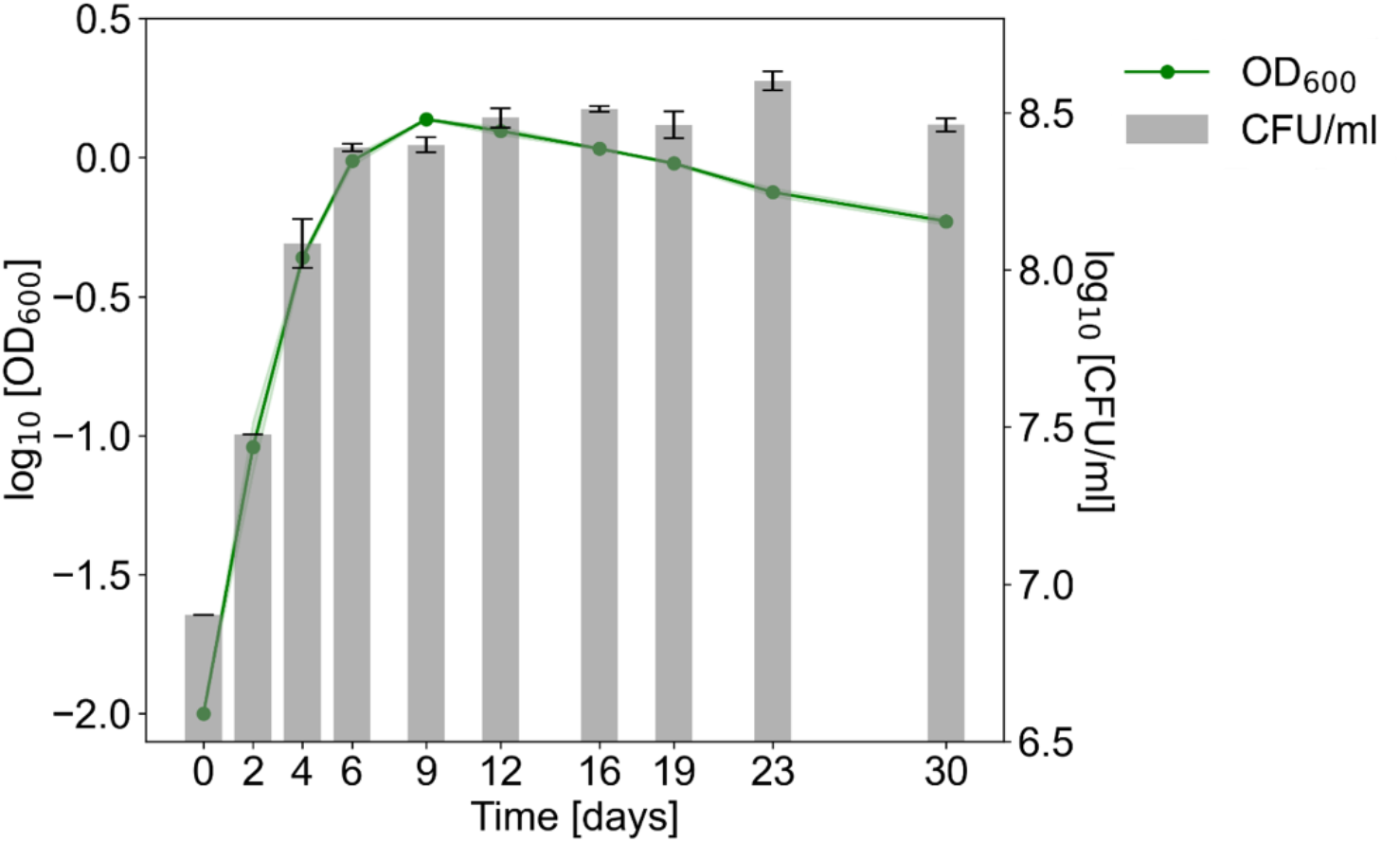
Growth dynamics of *P. inhibens* bacteria during starvation. Growth of *P. inhibens* on CNS medium with glucose as a carbon source was monitored over 30 days using OD_600_ (line) and CFU count (bars). Log-transformed data is presented as the mean of two biological replicates. Shaded area and error bars indicate standard deviations.

To evaluate nutrient depletion over the 30-day incubation period, we assessed the ability of the spent medium to support bacterial growth. At selected time points, cultures were filtered to remove cells, and the resulting cell-free filtrate (spent medium) was inoculated with fresh bacteria. Growth of the inoculum indicated that the medium could still support growth, whereas lack of growth suggested nutrient depletion. Rescue experiments in the form of targeted nutrient addition were then performed to identify the limiting nutrient. The results indicate that only the filtrate from exponential-phase cultures (day 3) supported bacterial growth, whereas filtrates from all later time points did not (Fig. 2). Supplementation of the filtrates with glucose restored growth (Fig. 2), indicating that carbon was depleted from day 7 onward.

**Figure 2.**
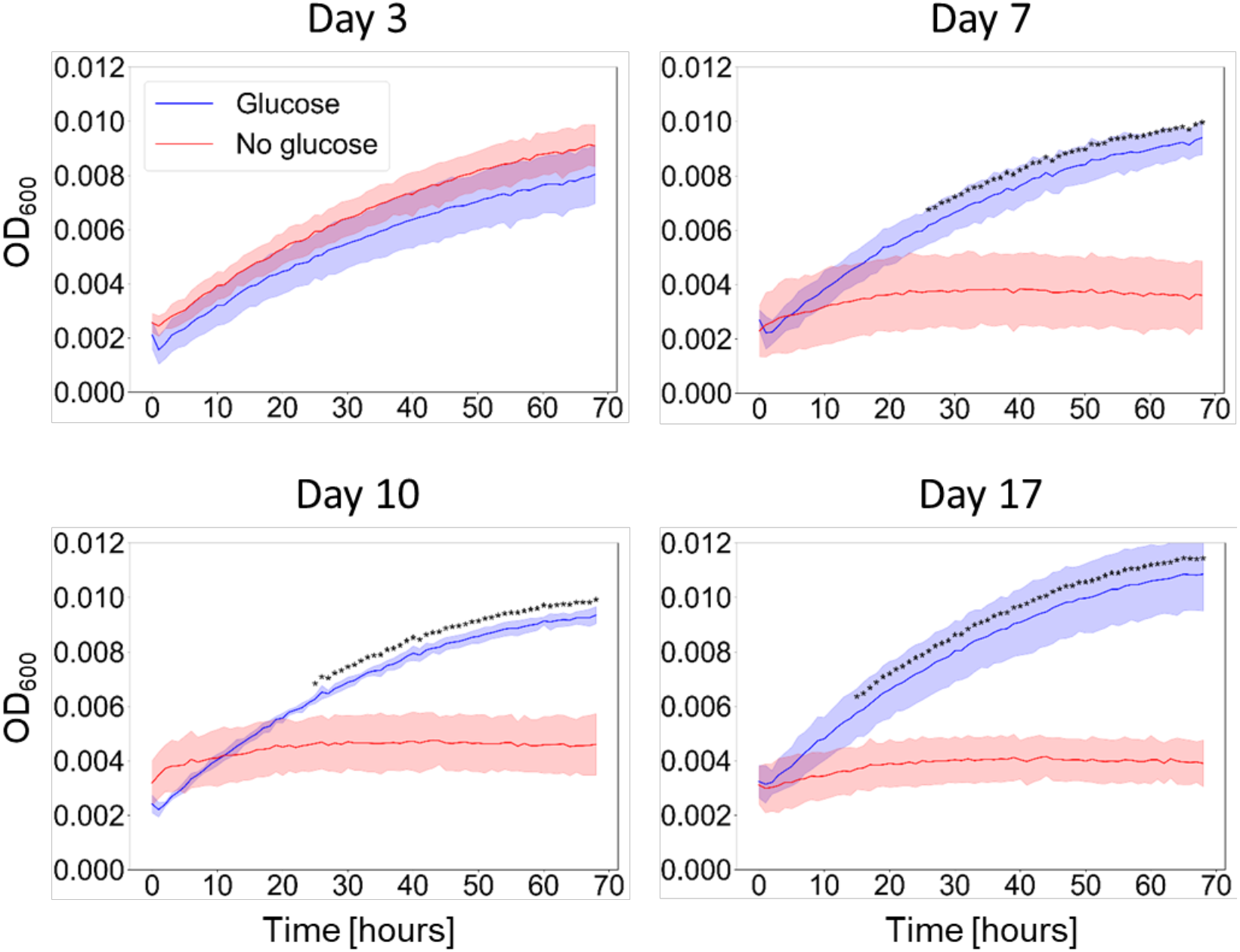
Carbon availability in filtrates of *P. inhibens* cultures during starvation. Bacterial cultures were grown in CNS medium for 3, 7, 10, or 17 days. The cultures were filtered through 0.22 µm filters to remove bacterial cells and obtain cell-free spent media. Fresh bacteria were inoculated into the spent media at an initial OD_600_ of 0.01 (red lines) or into the spent media supplemented with 6 mM glucose (blue lines). Bacterial growth was monitored over 68 hours by tracking OD_600_ (the culture age from whiuch the spent media is indicated above each panel). Lines represent the mean of three biological replicates, and shaded areas indicate the standard deviation. Asterisks indicate statistically significant differences between conditions with and without glucose supplementation, as determined by two-way ANOVA followed by Tukey’s post hoc test (p < 0.05).

### Cellular changes during starvation

Bacteria remained viable but did not grow during 30 days of incubation. To understand how cells survive in the absence of growth, we examined cellular morphology under carbon-depleted conditions. We first visualized live bacteria using phase-contrast microscopy to obtain an overview of starved cells. Our observations revealed that cell refractivity changed during growth and starvation; cells in early exponential phase (day 3) and late starvation (day 17) appeared darker, whereas cells in early stationary (day 7) and early starvation (day 10) phases were brighter (Fig. 3, upper panels).

**Figure 3.**
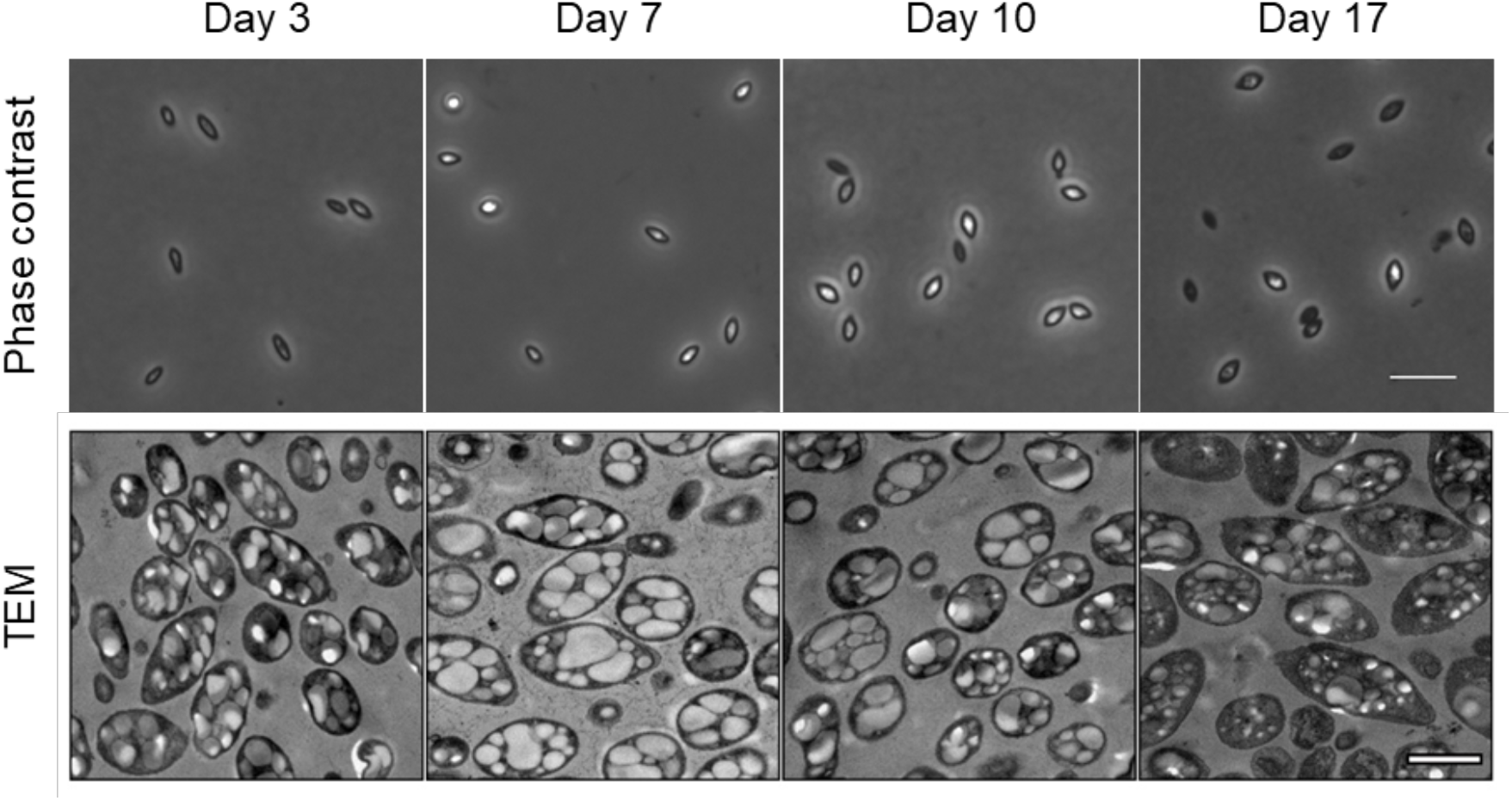
Morphology of starving *P. inhibens* bacteria. *P. inhibens* cells were sampled on days 3, 7, 10 and 17. Phase-contrast images show dark and bright cells, indicative of changing cell refractivity (upper panels, scale bar corresponds to 5 µm). Transmission Electron Microscopy (TEM) images show intracellular granules (lower panels, scale bar corresponds to 1 µm).

To identify intracellular structures that may contribute to these changes in refractivity, we analyzed subcellular morphological features at different growth and starvation stages using transmission electron microscopy (TEM). We collected bacterial cultures at representative time points and fixed all samples simultaneously (see Materials and Methods) prior to imaging. At all time points, the bacterial cells exhibited bright intracellular inclusions (Fig. 3, lower panels). The inclusions appeared to accumulate during exponential growth and seem to deplete during starvation. These inclusions resembled polyhydroxyalkanoate (PHA) carbon storage granules previously described in other bacterial species (Vandecandelaere et al. 2009; Xiao et al. 2015). The most common type of PHA is a polyhydroxybutyrate (PHB). To assess the capacity of *P. inhibens* to synthesize polyhydroxybutyrate (PHB), we analyzed its genome and identified the genes encoding the PHB biosynthetic pathway. This pathway converts acetyl-CoA to PHB through a three-step enzymatic process (Fig. S1). PHB biosynthesis involves three main genes: *phaA*, which encodes acetyl-CoA acetyltransferase; *phaB*, which encodes acetoacetyl-CoA reductase; and *phaC*, which encodes the PHB synthase (Fig. S1A). The *phaC* gene is essential for PHB production, while *phaA* and *phaB* may be functionally redundant (Koch and Forchhammer 2021).

### *P. inhibens* produces PHB storage granules

Although the granule morphology resembled that of PHB inclusions and genomic analysis confirmed the presence of the PHB biosynthetic pathway, we sought to verify that the observed intracellular granules are PHB storage granules. We therefore generated mutants lacking key genes in the PHB biosynthesis pathway (Fig. S1B) and compared these strains with wild-type (WT) bacteria with respect to granule morphology and composition.

We constructed the deletion strains Δ*phaA* (*phaA::Gm*^*R*^), Δ*phaB* (*phaB::Gm*^*R*^), and Δ*phaC* (*phaC::Km*^*R*^) and quantified the PHB content in the different strains using targeted metabolomics by liquid chromatography coupled to mass spectrometry (LC-MS). We hydrolyzed the samples and measured 3-hydroxybutyric acid, the PHB monomer released upon hydrolysis, against an external standard (see Materials and Methods). The Δ*phaA* and Δ*phaB* mutants showed significantly lower PHB concentrations than the WT, with approximately 2-fold and 10^2^-10^3^-fold reductions, respectively (Fig. 4A). In contrast, we detected only trace amounts of PHB in the Δ*phaC* strain (Table S1, File S1), suggesting that PHB production was severely impaired. Consistent with these results, TEM analysis of Δ*phaC* cells confirmed the absence of intracellular granules at all growth and starvation stages (Fig. 4B).

**Figure 4.**
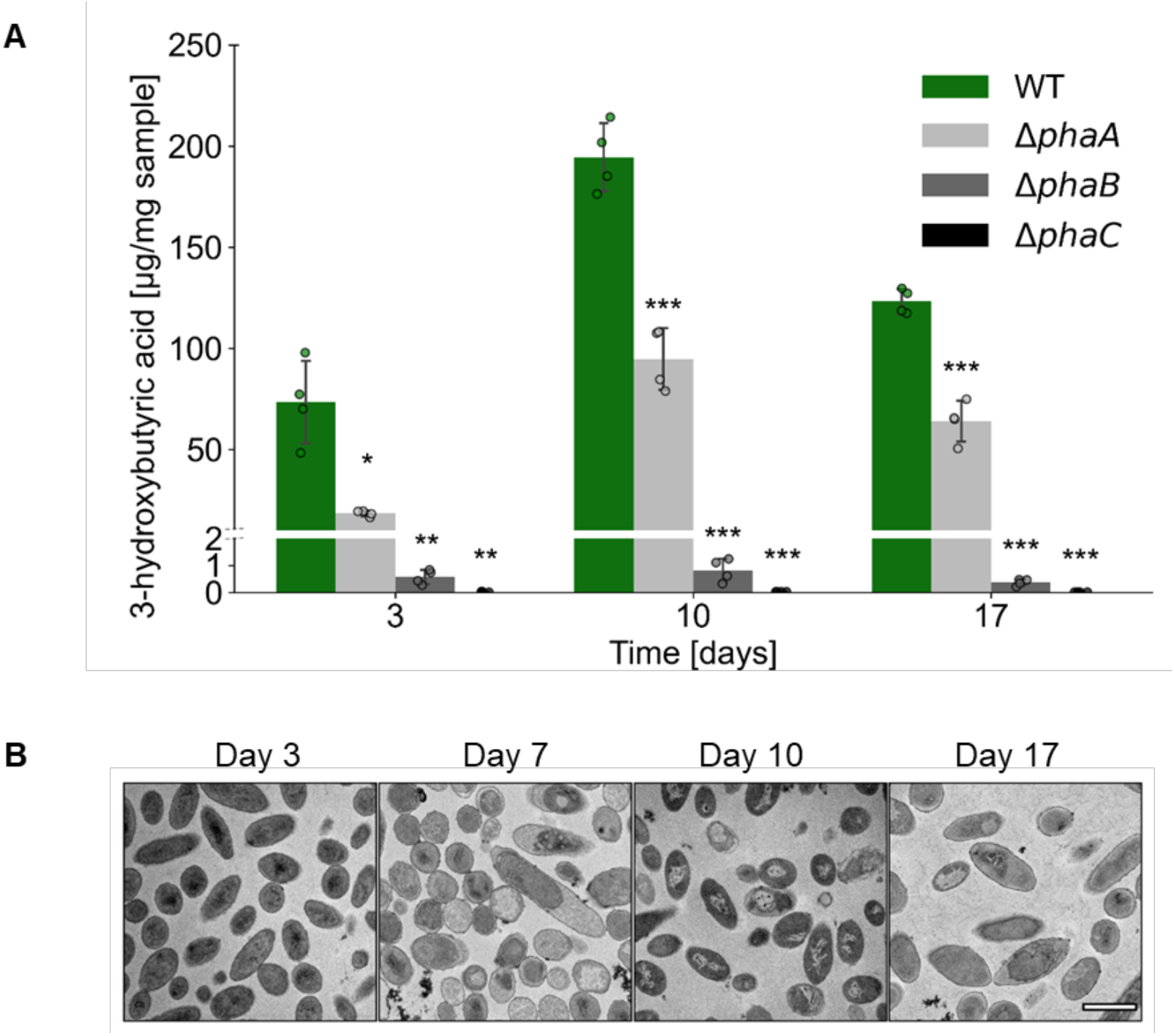
PHB accumulation in WT and mutant bacteria. **A**. Quantification of PHB was conducted by hydrolyzing the samples and measuring 3-hydroxybutyric acid, the monomer of PHB. Measurements were performed on wild-type (WT), Δ*phaA*, Δ*phaB*, and Δ*phaC* strains grown in CNS medium, using liquid chromatography coupled to mass spectrometry (LC-MS). Bars represent the mean concentration per milligram of sample from four biological replicates. Dots indicate individual replicate values, and error bars show the standard deviation. Comparisons between WT and each mutant strain were conducted at each time point using Welch’s *t*-tests, with *p*-values adjusted for multiple comparisons using the Benjamini–Hochberg false discovery rate (FDR) (*p* < 0.05, \**p* < 0.01, \*\**p* < 0.001). **B**. Transmission electron microscopy (TEM) images of Δ*phaC P. inhibens* cultures sampled on days 3, 7, 10, and 17. No intracellular inclusions (granules) were observed. Scale bar corresponds to 1 µm.

Overall, these results demonstrate that the intracellular granules observed in *P. inhibens* are PHB storage granules. Deletion of the *phaC* gene abolished PHB accumulation and granule formation, whereas deletion of the *phaA* or *phaB* genes reduced PHB production. These findings support redundant roles for PhaA and PhaB and a non-redundant, essential role for PhaC in PHB biosynthesis.

The *P. inhibens* WT strain exhibited temporal dynamics in PHB concentrations. PHB accumulated during the exponential phase (from day 3 to day 10) and was depleted during starvation (from day 10 to day 17). These trends are consistent with the granule dynamics observed by TEM (Fig. 3).

### PHB storage granules promote survival during starvation

*P. inhibens* cultures become progressively carbon-limited during starvation, and PHB granules decrease over time under these conditions. We therefore hypothesized that PHB storage granules support bacterial survival during carbon starvation. Previous studies proposed that PHB functions as a carbon reserve in marine bacteria under nutrient limitation (J. G. Wang and Bakken 1998; Malmcrona-Friberg et al. 1986; Jones and Rhodes-Roberts 1981; H. Wang et al. 2014; Kavitha et al. 2018), but a causal relationship was not yet established.

To test whether PHB storage contributes to starvation survival, we compared the viability of WT and mutant strains with reduced PHB content under starvation conditions. We inoculated cultures with comparable CFU counts and grew them in CNS medium (see Materials and Methods). We monitored growth during exponential and starvation phases by CFU enumeration, as OD_600_ measurements can be affected by optical differences between cells with and without granules.

All mutants exhibited growth comparable to the WT during the first three days of cultivation (Fig. 5). However, by the end of the starvation experiment (day 31), all mutants showed significantly lower cell counts than the WT. The Δ*phaA* mutant, which displayed a 2-fold reduction in PHB content, showed a decrease of approximately half an order of magnitude relative to WT on day 31. The Δ*phaB* and Δ*phaC* mutants, which had the lowest PHB levels (10^2^-10^3^-fold reduction or trace amounts, respectively), exhibited the most pronounced decline during starvation and reached cell counts approximately one order of magnitude lower than the WT on day 31. These results reveal a correlation between PHB content and survival under carbon limitation.

**Figure 5.**
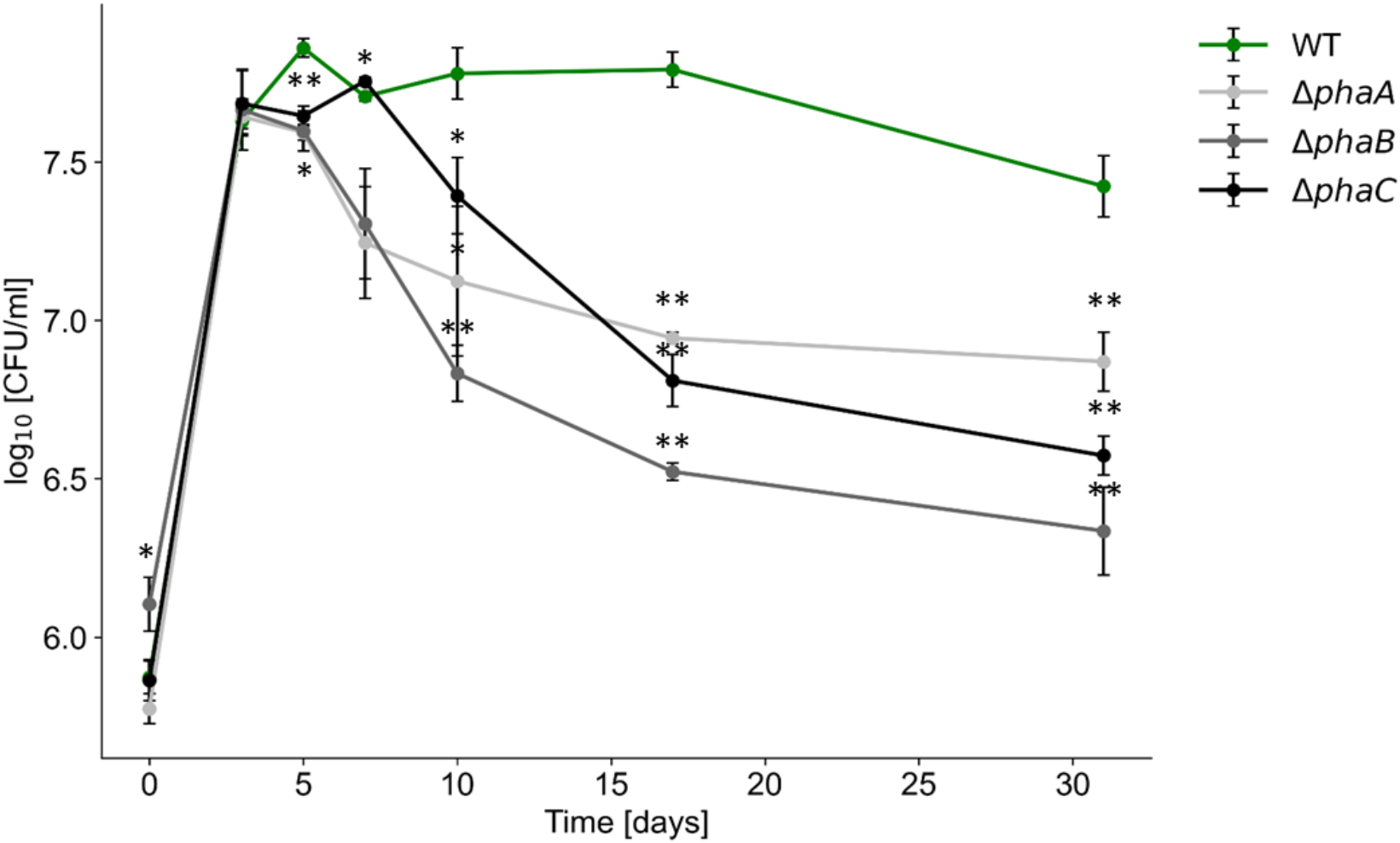
Deletion of PHB biosynthesis genes impairs *P. inhibens* survival under starvation. Viability of wild-type (WT) and Δ*phaA*, Δ*phaB, and* Δ*phaC* strains grown in CNS medium over 31 days was monitored by CFU counts. Each data point represents the mean of three biological replicates, and error bars indicate the standard deviation. Asterisks denote statistically significant differences between the WT and the corresponding mutant at a given time point, as determined by Welch’s *t*-tests with *p*-values adjusted for multiple comparisons using the Benjamini–Hochberg false discovery rate (FDR) (\**p* < 0.05, \*\**p* < 0.005).

Together, the findings highlight a role for PHB storage granules in supporting starvation survival and suggest that PHB accumulation represents an important strategy in marine heterotrophic bacteria. Notably, the Δ*phaC* strain, which lacks PHB granules, maintained viable counts above 10^6^ CFU/mL after 31 days of cultivation, indicating that additional mechanisms contribute to long-term survival under starvation.

### PHB granule accumulation may represent a common strategy among Roseobacters

PHB granule accumulation appears to support starvation survival in *P. inhibens*. We next asked whether this strategy is common among marine bacteria. Because *phaC* encodes the key enzyme in PHB biosynthesis, we analyzed the distribution of *phaC* orthologs within the Roseobacter group, the marine lineage of the *Roseobacteraceae* family to which *P. inhibens* belongs (see Materials and Methods). All examined members of the Roseobacter group encoded one to three copies of *phaC*, which clustered into distinct orthologous groups (File S2). This broad distribution indicates that the genetic capacity for PHB synthesis is widespread and conserved within the Roseobacter lineage.

To assess whether *phaC* is actively expressed in natural marine environments, we analyzed publicly available prokaryotic metatranscriptomic data from the TARA Oceans Atlas (see Materials and Methods). Among prokaryotic taxa expressing *phaC*, 25% belonged to the order *Rhodobacterales* (Fig. S2), which includes the *Roseobacteraceae* family. Most transcripts were affiliated with bacteria, including members of the α-, β-, and γ-proteobacteria, whereas a smaller fraction was assigned to archaea, primarily Thaumarchaeota.

Finally, we examined whether bacteria outside the Roseobacter lineage can survive starvation and whether the presence of a *phaC* ortholog predicts this capacity. To this end, we performed starvation experiments with diverse marine bacterial species that either encode a *phaC* ortholog (*Marinobacter* sp., *Qipengyuania flava*, and *Oceanicaulis alexandrii*) or lack it (*Alteromonas macleodii, Balneola vulgaris*, and *Ulvibacter litoralis*), representing multiple phylogenetic classes (Fig. S3A, Table 1). We included *P. inhibens* WT and Δ*phaC* strains as controls.

**Table 1.** Marine bacterial species from four taxonomic classes used in the starvation experiment (shown in Fig. S3) and their reference genomes.

| Species | Class | *phaC* ortholog | Reference genome |
| --- | --- | --- | --- |
| *Phaeobacter inhibens* DSM 17395 | *Alphaproteobacteria* | Yes | GCF_000154765.2 |
| *Phaeobacter inhibens* DSM 17395 Δ*phaC* | *Alphaproteobacteria* | No | This study |
| *Marinobacter* sp. | *Gammaproteobacteria* | Yes | https://doi.org/10.5281/zenodo.7520020 |
| *Alteromonas macleodii* | *Gammaproteobacteria* | No | https://doi.org/10.5281/zenodo.7519869 |
| *Qipengyuania flava* (previously known as *Erytherobacter flavus*) | *Alphaproteobacteria* | Yes | https://doi.org/10.5281/zenodo.7520027 |
| *Oceanicaulis alexandrii* DSM 11625 | *Alphaproteobacteria* | Yes | GCF_000420265.1 |
| *Balneola vulgaris* DSM 17893 | *Balneolia* | No | GCF_000375465.1 |
| *Ulviabacter litoralis* DSM 16195 | *Flavobacteriia* | No | GCF_900102055.1 |

The different species exhibited varying starvation survival capabilities. We did not observe a clear correlation between the presence of *phaC* and the ability to survive starvation (Fig. S3B). These results indicate that additional, yet unidentified mechanisms contribute to starvation survival in marine bacteria.

## Discussion

### PHB storage promotes survival during carbon starvation

Marine bacteria commonly store carbon in intracellular granules, including polyhydroxybutyrate (PHB), which is widespread in marine environments (H. M. Alvarez et al. 1997). Although PHB has attracted attention for bioplastic production, its ecological role remains unclear (López-Cortés et al. 2008; Mohanrasu et al. 2018; Kavitha et al. 2018). Previous studies showed PHB depletion in the marine strains *Vibrio sp*. and *Sphaerotilus discophorus* during starvation, suggesting that PHB serves as a carbon and energy reserve (Mårdén et al. 1985; Stokes and Parson 1968).

We used a genetic approach to test whether PHB directly supports bacterial survival during carbon starvation. Deleting the *phaC* gene in *P. inhibens* eliminated storage granules and reduced survival under starvation compared to the WT. We still detected trace amounts of 3-hydroxybutyric acid, a PHB monomer, in the Δ*phaC* mutant, suggesting involvement in other metabolic pathways. During starvation, *P. inhibens* WT maintained stable CFU counts, while OD_600_ values gradually decreased (Fig. 1). This divergence between viability and optical density likely reflects the progressive consumption of PHB granules observed here, together with a reduction in cell size that has been previously reported in starving bacteria (Amy and Morita 1983; Bergkessel and Delavaine 2021). Both processes would reduce OD_600_ without affecting CFU counts.

### PHB enables marine bacteria to cope with fluctuating marine environment

Carbon storage granules help marine bacteria adapt to fluctuating environmental conditions. In the ocean, nutrient pulses create feast-and-famine dynamics that shape bacterial physiology (Moriarty and Bell 1993). PHB acts as a metabolic buffer that promotes survival during scarcity and supports rapid growth during periods of nutrient availability. PHB may also increase bacterial resistance to environmental stress, including thermal, oxidative, and osmotic challenges (Müller-Santos et al. 2021; Sedlacek et al. 2019).

Fluctuating environments can also generate phenotypic heterogeneity within bacterial populations (Morawska et al. 2022). Because we measured PHB content and cell viability at the population level, we may overlook variation between individual cells. Microscopy phase-contrast images (Fig. 3) from days 10 and 17 reveal cells with different refractive properties, suggesting heterogeneity in intracellular PHB content. Such variability may represent a bet-hedging strategy that enhances survival under changing conditions.

### Diverse survival strategies across marine bacteria

Although the Δ*phaC* mutant showed reduced survival, it still persisted during prolonged starvation. This result indicates that PHB contributes to survival but is not the only mechanism involved. Previous work in soil bacteria has shown that PHB enhances starvation survival within strains but that survival remains species-specific (J. G. Wang and Bakken 1998). Consistent with this, several marine strains tested here lack the capacity to synthesize PHB, yet survived starvation (Fig. S3B). These findings suggest that marine bacteria use multiple strategies to survive under carbon limitation. These can include metabolic remodeling and reduction in cell size (Bergkessel and Delavaine 2021), scavenging of trace nutrients (Mårdén et al. 1987; Davis and Robb 1985), recycling of cellular components or dead cells (S. J. Schink et al. 2019; Takano et al. 2017), and the use of alternative storage compounds. Indeed, marine bacteria can accumulate other carbon reserves such as wax esters and triacylglycerols (H. M. Alvarez et al. 1997; Héctor M. Alvarez 2016), and polyphosphate as a phosphorus storage compound (Gao et al. 2025). The uneven taxonomic distribution of the *phaC* gene, coupled with its lack of strict correlation with survival, suggests that bacterial lineages rely on diverse and potentially complementary strategies to survive starvation.

### Ecological implications of storage granules for algal-associated bacteria

Heterotrophic marine bacteria rely on algal partners as a major source of organic carbon. The algal carbon supply fluctuates over time between bloom events and across diel cycles (Gasol et al. 2016; Granum et al. 2002; Doblin et al. 2011). Consequently, bacteria must endure repeated periods of carbon limitation. Intracellular storage compounds such as PHB can support survival between nutrient pulses. By supporting survival during carbon limitation, PHB storage may help maintain stable bacterial populations associated with algae. When nutrients become available, these reserves enable rapid growth, promoting population recovery and influencing bacterial succession during bloom progression (Zhu et al. 2023; Sison-Mangus et al. 2016).

Our ability to detect bacterial starvation in the ocean remains limited, and we still lack a clear picture of how common starvation is and how bacteria survive it. In this study, we show that PHB storage directly supports survival during carbon starvation, providing a concrete example of a molecular mechanism that links physiology to ecological function. As we identify more genes, metabolites, and pathways involved in starvation survival, we can use this knowledge to analyze environmental datasets and detect signatures of starvation in natural microbial communities. Each new mechanism expands the set of molecular markers that can be tracked in the ocean. By combining laboratory discoveries with environmental data, we can begin to map when and how bacteria experience starvation, and better understand how these survival strategies shape microbial community dynamics and carbon cycling in marine ecosystems.

## Materials and Methods

### Bacterial strains and general culture conditions

The bacterial strain *Phaeobacter inhibens* DSM 17395 was purchased from the German Collection of Microorganisms and Cell Cultures (DSMZ, Braunschweig, Germany). Bacteria were plated from 20% glycerol stocks (stored at -80°C) onto ½ YTSS agar plates (2 g yeast extract, Sigma-Aldrich, USA; 1.25 g tryptone, Sigma-Aldrich, USA; 20 g sea salts, Sigma-Aldrich, USA; and 16 g agar per liter, BD Difco, USA). After 48 hours of incubation at 30°C, single colonies were inoculated into liquid CNS medium to establish a pre-culture. CNS was prepared with 6 mM glucose (Sigma-Aldrich, USA), 5 mM NH_4_Cl (Sigma-Aldrich, USA), 33 mM Na_2_SO_4_ (Fisher Scientific, UK), and diluted in artificial sea water (ASW). The ASW medium was based on a previously published protocol (Goyet and Poisson 1989) and contained mineral salts (NaCl, 409.41 mM; Na_2_SO_4_, 28.22 mM; KCl, 9.08 mM; KBr, 0.82 mM; NaF, 0.07 mM; Na_2_CO_3_, 0.20 mM; NaHCO_3_, 2 mM; MgCl·6H_2_O, 50.66 mM; CaCl_2_, 10.2 mM; SrCl_2_·6H_2_O, 0.09 mM), f/2 vitamins (thiamine HCl, 100 μg l^−1^; biotin, 0.5 μg l^−1^; vitamin B_12_, 0.5 μg l^−1^), L1 trace elements (Na_2_EDTA·2H_2_O, 4.36 mg l^−1^; FeCl_3_·6H_2_O, 3.15 mg l^−1^; MnCl_2_·4H_2_O, 178.1 μg l^−1^; ZnSO_4_·7H_2_O, 23 μg l^−1^; CoCl_2_·6H_2_O, 11.9 μg l^−1^; CuSO_4_·5H_2_O, 2.5 μg l^−1^; Na_2_MoO_4_·2H_2_O, 19.9 μg l^−1^; H_2_SeO_3_, 1.29 μg l^−1^; NiSO_4_·6H_2_O, 2.63 μg l^−1^; Na_3_VO_4_, 1.84 μg l^−1^; K_2_CrO_4_, 1.94 μg l^−1^) and L1 nutrients (NaNO_3_, 882 μM; NaH_2_PO_4_·2H_2_O, 36.22 μM). All components were dissolved in Milli-Q water (IQ 7003; Merck) and the pH was adjusted to 8.2 using HCl. Stock solutions of f/2 vitamins, L1 trace elements, and L1 nutrients were purchased from the Bigelow Laboratory for Ocean Sciences (Boothbay, ME, USA). Pre-cultures were grown in 30°C in the dark with shaking at 130 rpm for at least 48 hours to reach stationary phase.

The mutant strains *Phaeobacter inhibens* DSM 17395 Δ*phaA* (*phaA*::Gm^R^), Δ*phaB* (*phaB*::Gm^R^), and Δ*phaC* (*phaC*::Km^R^) were generated in this study (see below) and cultivated under the same conditions as the wild-type (WT) *P. inhibens* DSM 17395 strain. Antibiotics were added as appropriate: 30 μg/ml gentamicin or 150 μg/ml kanamycin to both agar plates and liquid media for pre-cultures.

The bacterial strains *Oceanicaulis alexandrii* DSM 11625, *Balneola vulgaris* DSM 17893 and *Ulvibacter litoralis* DSM 16195 were obtained from the German Collection of Microorganisms and Cell Cultures (DSMZ, Braunschweig, Germany). Strains *Marinobacter* sp., *Alteromonas macleodii* and *Qipengyuania flava* (previously known as *Erytherobacter flavus* (Lee and Kim 2020)) were previously isolated from *Emiliania huxleyi* CCMP1516 and sequenced (Beiralas et al. 2023). All bacteria were grown in Marine Broth 2216 medium (MB) (Difco, USA) in liquid cultures or on agar plates under the cultivation conditions described above.

### Deletion of *phaA, phaB* and *phaC* genes in *P. inhibens*

To generate the *phaA, phaB*, and *phaC* gene deletion mutants of *Phaeobacter inhibens* DSM 17395, the genes PGA1_RS01675, PGA1_RS01670, and PGA1_RS10360 were replaced with antibiotic resistance cassettes, resulting in the mutant strains ES320, ES319, and ES321, respectively. A gentamicin resistance cassette was used to replace the *phaA* and *phaB* genes, while a kanamycin resistance cassette was used to replace the *phaC* gene. Regions of approximately 1000 bp upstream and downstream of each gene were PCR-amplified using primers 1323-1334 (Table 2). The amplified fragments were assembled and cloned either into the pDN18 vector for constructs containing the gentamicin resistance cassette, or into the pCR8/GW/TOPO vector with the resistance cassette for the construct containing the kanamycin resistance cassette (see Table 3) using restriction-free cloning (Unger et al. 2010). The resulting plasmids pLK1, pLK2 and pYT1 for *phaA, phaB* and *phaC* genes deletion respectively (Table 3) were introduced into competent *P. inhibens* bacteria by electroporation. Preparation of competent cells was performed as previously described (Piekarski et al. 2009), with slight modifications. Briefly, cells were grown to an OD_600_ of 0.6 in ½ YTSS with 40 g/L sea salts (also termed full salts medium). Bacteria were washed three times with 10% (v/v) ice-cold glycerol, centrifuged each time for 5 min at 4 °C and 6200 g. The competent cells were subsequently adjusted to an OD_600_ of 20 using 10% (v/v) ice-cold glycerol, frozen in liquid nitrogen and stored at -80°C. Electroporation was conducted with 300 μl aliquots of electrocompetent cells, using 10 µg of DNA in a 2 mm cuvette at 2.5 V, followed by 4 h recovery with shaking in ½ YTSS with full sea-salts concentration. The transformed cells were then plated on ½ YTSS plates containing 30 μg/ml gentamicin (Δ*phaA* and Δ*phaB*) or 150 μg/ml kanamycin (Δ*phaC*), and resistant colonies were validated by PCR (primers 1335-1336 and 1339-1342; see Table 2) and DNA sequencing.

**Table 2.**
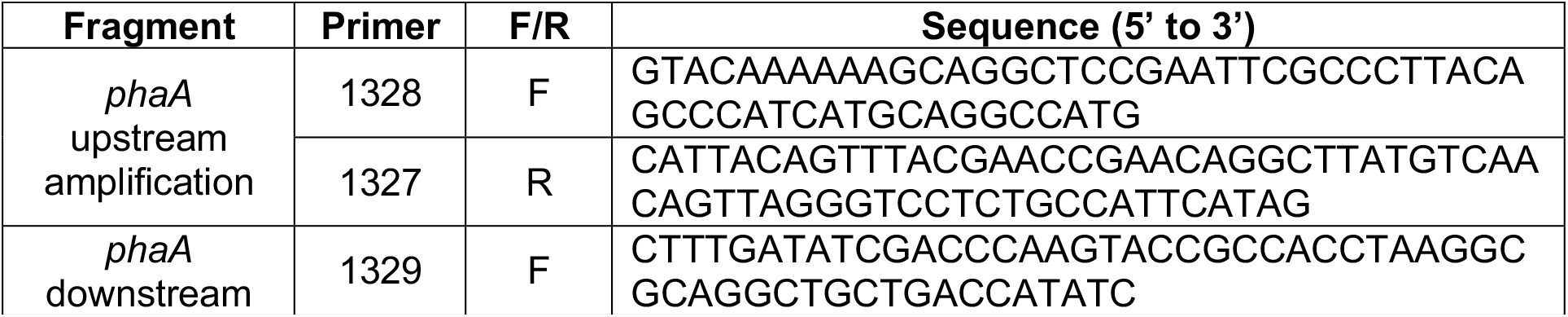
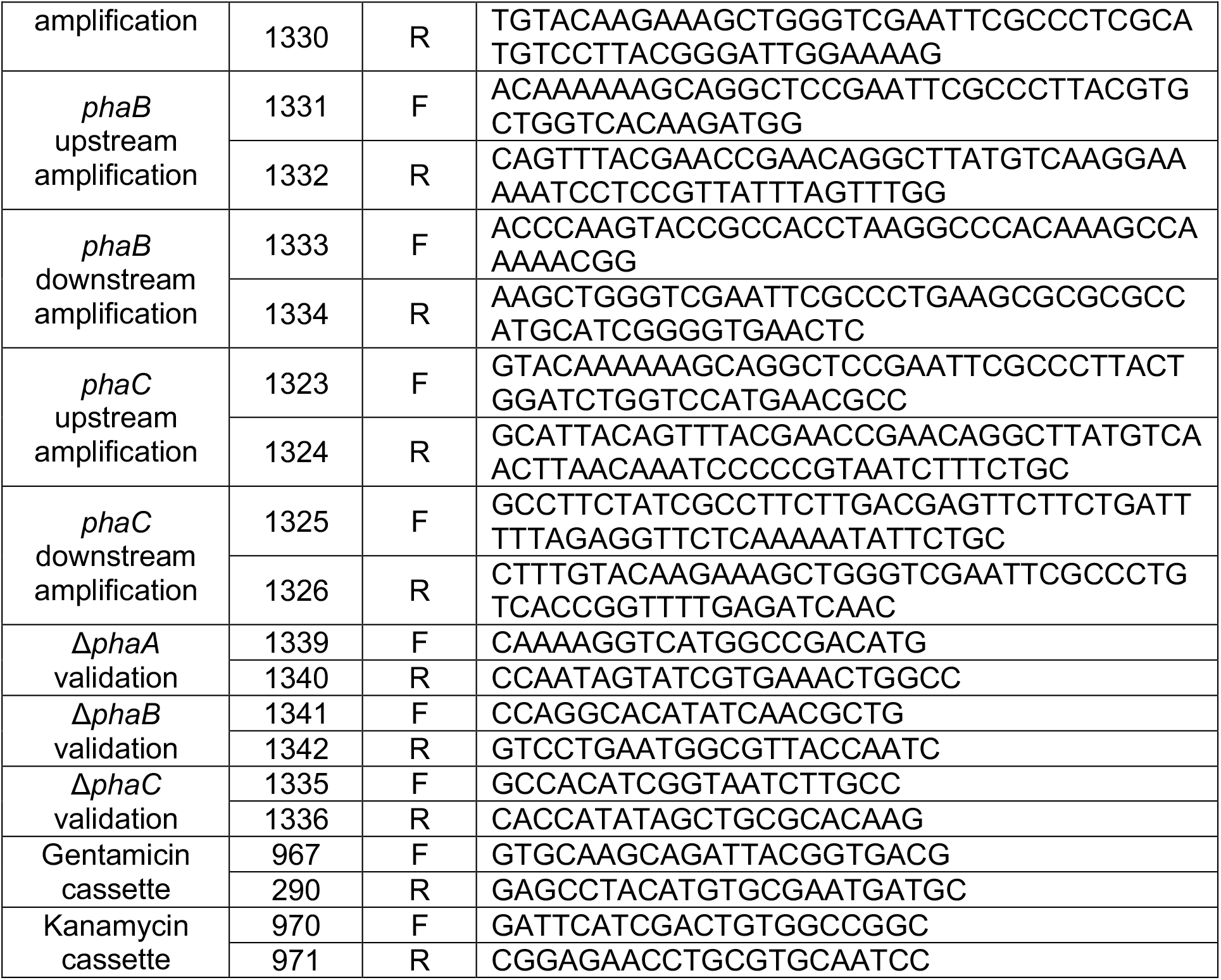
Primers used in this study.

**Table 3.** Plasmids used in this study.

| Name | Construct | Origin | Comments |
| --- | --- | --- | --- |
| pLK1 | *phaA*::Gm-TOPO8 | Current study | Constructed using DN18 for *phaA* gene replacement with the gentamicin resistance cassette |
| pLK2 | *phaB*::Gm-TOPO8 | Current study | Constructed using DN18 for *phaB* gene replacement with the gentamicin resistance cassette |
| pYT1 | *phaC*::Km-TOPO8 | Current study | Constructed using pCR8/GW/TOPO vector (Invitrogen) and kanamycin resistance cassette for *phaC* gene replacement |
| pDN18 | Gm-TOPO8 | Lab collection | Constructed using pCR8/GW/TOPO vector (Invitrogen), and used in the current study as a vector carrying the gentamicin resistance cassette |

Once the mutant strains were generated, cultures were grown without antibiotics. To verify the absence of wild-type (WT) contamination, samples were plated on agar plates both with and without antibiotics, yielding identical colony counts. Additionally, PCR amplification using the primers 290, 267, 970, 971, 1335, 1336, 1339-1342 (Table 2) confirmed the absence of WT cells.

### Culture preparation and monitoring bacterial growth

For all experiments, the bacterial concentrations in pre-cultures were determined by measuring OD_600_ values using an Ultrospec 2100 pro spectrophotometer (Biochrom, Cambridge, UK) with plastic cuvettes. Cultures were then inoculated into CNS or MB media to an OD_600_ of 0.01. Cultures of 100 ml with two biological replicates were used to generate the long-term growth curve (Fig. 1) and cultures of 30 ml with at least three biological replicates were used for all other experiments. All cultures were maintained in standing conditions within a growth room at a temperature of 18 °C under a light/dark cycle of 16/8 h. Light intensity during the illuminated period was maintained at 150 µmoles/m^2^/s. Cultures of mutant strains were cultivated without antibiotic addition to ensure similar growth conditions to WT cultures. Cultures were sampled to ensure the absence of WT contamination using both plating and PCR, as described above. Bacterial growth in cultures was monitored by OD_600_ measurements and by colony-forming unit (CFU) counts on agar plates. For OD_600_ evaluation, 800 μl of diluted culture was measured in 1 cm cuvettes. For CFU count, bacterial cultures were serially diluted in sterile ASW and plated on corresponding agar plates: ½ YTSS for *P. inhibens* WT, ½ YTSS with 30 μg/ml gentamicin for Δ*phaA* and Δ*phaB* mutants, and ½ YTSS with 150 μg/ml kanamycin for Δ*phaC* mutant. For starvation experiment with additional bacterial strains (Fig. S3B), all bacterial cultures were plated on MB agar plates instead. After two days of incubation at 30°C, the number of colonies was evaluated, and corresponding CFU/ml values were calculated.

### Filtrate media experiment

*P. inhibens* WT cultures were grown in biological triplicates in CNS medium for 3, 7, 10 or 17 days. The growth media were collected on the same day and filtered through 0.22 µm filters to remove bacterial cells. Fresh *P. inhibens* WT cultures (grown in CNS) were inoculated at an initial OD_600_ of 0.01 into the filtered media or filtered media supplemented with 6 mM glucose (150 μl per each well) in triplicates in a 96-well plate. Corresponding media without bacterial addition were used as a blank control. Each well was overlaid with 50 µl hexadecane to prevent evaporation (Sklodowska and Jakiela 2017). Bacterial growth was monitored using an Infinite 200 Pro M Plex plate reader (Tecan Group Ltd., Männedorf, Switzerland) with alternating cycles of measurements and 1 hour of incubation at 30°C. Optical density measurements were conducted at 600 nm following the shaking step. Corresponding blank values were subtracted from each data point before the analysis.

### Light microscopy

Phase contrast images were obtained using a Nikon Eclipse Ti2-E inverted microscope equipped with a CFI Plan Apochromat DM 100X objective lens (Nikon, Tokyo, Japan). All samples were spotted on thin 1% agarose pads for visualization at room temperature. Images were acquired using an Andor Zyla 4.2 camera controlled by NIS-Elements software. Images were processed identically.

### Transmission electron microscopy

For transmission electron microscopy (TEM) visualization, 30 ml cultures of 3, 7, 10 and 17 days of *P. inhibens* WT and Δ*phaC* mutant were prepared in an asynchronous manner to ensure biomass collection on the same day.

All wild-type and mutant cells were fixed overnight with a solution of 2% glutaraldehyde, 4% paraformaldehyde in artificial seawater (ASW) buffer. Cells were rinsed with ASW, pelleted via centrifugation, and embedded in a 3.4% agarose solution which was then hardened on ice to prevent dispersion of cells during the secondary fixation and embedding process. Agarose-embedded cell pellets were fixed and stained for one hour with an osmium tetroxide solution (1% OsO4, 0.5% potassium dichromate, 0.5% potassium hexacyanoferrate in ASW), rinsed with ASW followed by milli-Q H_2_O, and then stained for another hour in 2% uranyl acetate. Pellets were rinsed again with milli-Q H_2_0 and then dehydrated with a series of ethanol dilutions (50%, 70%, 96%, 100%) and then transferred to 100% acetone. Pellets were then embedded in epon (EMS, PA, USA), diluted with anhydrous acetone in increasing concentrations: 25%, 50, 75%, and 100%. Cell pellets were finally embedded in 100% epon within rectangular molds, dried at 60°C for 24 hours, and stored until sectioning. Sample blocks were sectioned to a thickness of 70 nm with an EM UC7 ultramicrotome (Leica microsystems, Vienna, Austria) equipped with an Ultra 45° diamond knife (Diatome Ltd, Nidau, Switzerland) and were mounted on formvar and carbon-coated 200 mesh copper grids (EMS). Sections were stained with Reynolds lead citrate and imaged at room temperature with 120 kV using a Tecnai Spirit (Bio Twin) microscope transmission electron microscope (Thermo Fisher Scientific, The Netherlands) equipped with a bottom-mounted Gatan OneView 4k x 4k CMOS camera.

### Detection of polyhydroxybutyrate (PHB) by mass-spectrometry

Intracellular polyhydroxybutyrate content was estimated based on the monomer 3-hydroxybutyric acid. Sample preparation was based on (Chen and Yu 2005). Pellets (30 mg) were lyophilized, then 300 µl KOH 4N was added, followed by 10 min sonication and hydrolysis for 2 hours at 70°C in a shaker at 500 rpm. After cooling to room temperature, pH was neutralized by adding 300 µl HCl 4N and adjusted with KOH or HCl. Samples were centrifuged and transferred to vials. Dilution of 10, 100 or 1000 times in MeOH : DDW 1:1 and reinjection were performed if needed, according to concentration of 3-hydroxybutyric acid.

The measurement of 3-hydroxybutyric acid was done as described at (Zheng et al. 2015) with minor modifications detailed below. Briefly, analysis was performed using UPLC-ESI-MS/MS equipped with Acquity UPLC I class system (Waters). The LC separation was done using the Atlantis Premier BEH Z-HILIC (100 mm × 2.1 mm) (Waters). Mobile phase B consisted of acetonitrile, and mobile phase A consisted of 20 mM ammonium carbonate with 0.1% ammonium hydroxide in water–acetonitrile mixture (80:20, v/v). The flow rate was maintained at 200 µL min^−1^, and the gradient was as follows: 0-2 min, 75% B; 14 min, 25% B; 18 min, 25% B; 19 min, 75% B, held for 4 min. Column temperature was set to 45°C and injection volume was 0.5 µl. MS detector (Waters TQ-XS) was equipped with ESI source. The measurement was performed in negative ionization mode. The source and de-solvation temperatures were maintained at 150°C and 600°C, respectively. The capillary voltage was set to 1.0 kV, and cone voltage to 20 V. Nitrogen was used as de-solvation gas and cone gas at the flow rate of 1000 L×h^-1^ and 150 L×h^-1^, respectively. Quantification of 3-hydroxybutyric acid was performed using MRM 102.9 > 58.9 (Collision energy 10 eV), and the second MRM 102.9 > 40.8 (Collision energy 20 eV) was used for identification. SIR of mass 85.0 was used for detection of crotonic acid (putative identification, without standard). Data were processed with the MassLynx software with Targetlynx (Waters). Quantification of 3-hydroxybutyric acid in samples was performed against an external calibration curve prepared of 3-hydroxybutyric acid (Aldrich cat 166898-1G) in MeOH :DDW (1:1, v/v).

### Bioinformatical analysis of the *phaC* gene orthologous in Roseobacters

The protein fasta files of each of the samples were downloaded and used as input in OrthoFinder (Emms and Kelly 2019) to identify orthologous relationships among gene sequences of the different species. OrthoFinder was run with the default inflation value of 1.5.

Out of the 19,325 orthogroups detected, 107 had a single copy in all the analyzed genomes. These orthogroups were used to reconstruct a species tree. The sequences of each orthogroup were aligned separately using the l-ins-i algorithm in MAFFT (Katoh and Standley 2013) and the alignment was trimmed with trimAl (Capella-Gutiérrez et al. 2009) to exclude poorly aligned regions. The trimmed alignments were then concatenated and a tree was reconstructed using RAxML 8 (Stamatakis 2014), allowing automatic detection of model evolution for each sequence partition, and using 100 replicates of rapid bootstrap analysis for branch support.

Orthogroups with sequences annotated as PHB Polymerase or as PhaC were included in a single file, and a tree was reconstructed as described above, in a single partition analysis. Sequences from the same sample, but from different orthogroups, were kept separate. The trees were manually inspected using ete3 (Huerta-Cepas et al. 2016) to identify polyphyletic genera and misidentifications, and to confirm and refine the orthological relationships among the gene sequences. PhaC paralogs IDs were assigned to top level monophyletic clusters in the gene tree.

In the gene tree, which should be regarded as unrooted, two large and relatively homogeneous clusters were denoted as paralogs P1 and P2. An additional cluster was denoted U, with a subdivision to U1, U2 and U3. Cluster P2 was subdivided into P2.1, P2.2, and P2.3 to capture the three monophyletic clusters comprising it. A large monophyletic cluster within P1 was denoted P1.1, with the remaining leaves in the cluster comprising a paraphyletic group.

### Environmental data

For environmental data analysis, we used publicly available metatranscriptomic data from the Ocean Gene Atlas (Vernette et al. 2022; Villar et al. 2018). TARA Ocean Microbiome Reference Gene Catalog v2 was used as a reference dataset, and the protein sequence of *P. inhibens* PhaC protein (WP_014880410.1) was submitted as entry sequence. Blastp searches were performed using standard settings.

### Phylogenetic tree reconstruction for used marine strains

To demonstrate the phylogenetic relationships between species used in this study (Fig. S3), we reconstructed the phylogenetic tree using phyloT with the following NCBI taxa: 28108, 1121104, 192812, 391619, 1122613, 227084 and 50741. Standard parameters were used (expanded internal nodes, polytomy), and Interactive Tree Of Life (iTOL) (Letunic and Bork 2021) was used for visualization.

### Statistics

At least three biological replicates were used for each treatment, except for long-term cultures (Fig. 1) where two biological replicates were used. Mean of all technical replicates was used as representative of a corresponding biological replicate. Growth data from Figures 1, 5 and S3 were first log_10_-transformed, and then mean and standard deviation were calculated. For statistical analysis, Welch’s t-tests with p-values adjusted for multiple comparisons (the Benjamini–Hochberg false discovery rate) were used to compare WT to mutant strains. Two-way ANOVA followed by Tukey’s post hoc test was used for comparisons between multiple treatments. Statistical tests were performed using Spyder 6.0.1 (Python Software Foundation).

## Acknowledgements

We thank Prof. Assaf Vardi and Dr. Constanze Kuhlisch for sharing bacterial strains. We are grateful to all members of the Segev lab for valuable discussions and input.

## Funding

O.S. received the Institute for Environmental Sustainability Fellowship. The study was funded by the Israel Science Foundation (ISF 692/24), the European Research Council (ERC StG 101075514) and the de Botton Center for Marine Science, granted to E.S.

## Competing interests

The authors declare no competing interests.

## Notes

### Competing Interest Statement

The authors have declared no competing interest.

