## Supplementary material for "Intracellular carbon storage enables starvation survival in marine bacteria": File S2

| | P1 | P1.1 | P2.1 | P2.2 | U1 | U2 | U3 | Ref |
|---|---|---|---|---|---|---|---|---|
| **Species** | | | | | | | | |
| **Actibacterium_atlanticum__22II_S11_z10** | ● | | ● | | | | | WP_0352515641<br>WP_0818064561 |
| **Actibacterium_lipolyticum__CECT_8621** | ● | | ● | | | | | WP_0939674861<br>WP_0939680671 |
| **Actibacterium_mucosum__KCTC_23349** | ● | | | | | | | WP_0352621301 |
| **Actibacterium_naphthalenivorans__DSM_105040** | ● | | ● | | | | | WP_0545393751<br>WP_0545400951 |
| **Actibacterium_pelagium__JN33** | ● | | | | | | ● | WP_0955950231<br>WP_0955949901 |
| **Actibacterium_ureilyticum__LS_811** | ● | | ● | | | | | WP_0955905731<br>WP_0955874951 |
| **Algirhabdus_cladophorae__KMM_6494** | ● | | | | | | | WP_4208608291 |
| **Aliiruegeria_haliotis__DSM_29328** | ● | | ● | | | | | WP_1062051401<br>WP_1062028571 |
| **Aliiruegeria_lutimaris__DSM_25294** | ● | | ● | | | | | WP_0931481291<br>WP_2445207571 |
| **Aliiruegeria_sabulilitoris__GJMS_35** | ● | | ● | | | | | WP_0683088921<br>WP_2317010621 |
| **Aliisedimentitalea_scapharcae__KCTC_42119** | | ● | | | | | | WP_4066487121 |
| **Alloyangia_mangrovi__SAOS_153D** | | ● | ● | | | | | WP_0958829391<br>WP_2603490191 |
| **Alloyangia_pacifica__CGMCC_1.3455** | | ● | ● | | | | | WP_0924251931<br>WP_0924281031 |
| **Celeribacter_arenosi__JCM_17190** | ● | | | | | | | WP_3448469221 |
| **Celeribacter_baekdonensis__DSM_27375** | ● | | | ● | | | | WP_0746418111<br>WP_0746409241 |
| **Celeribacter_ethanolicus__TSPH2** | ● | | | | | | | WP_0968055281 |
| **Celeribacter_halophilus__DSM_26270** | ● | | | | | | | WP_0666031441 |
| **Celeribacter_indicus__P73** | ● | | | ● | | | | WP_0438706131<br>WP_0524532231 |
| **Celeribacter_litoreus__ASW11_22** | ● | | ● | | | | | WP_2245043481<br>WP_2245042901 |
| **Celeribacter_marinus__IMCC12053** | ● | | | | | | | WP_0622194681 |
| **Celeribacter_naphthalenivorans__EaN35_2** | ● | | | | | | | WP_2265530441 |
| **Celeribacter_neptunius__DSM_26471** | ● | | ● | | | | | WP_0900590381<br>WP_2181513111 |
| **Celeribacter_persicus__DSM_100434** | ● | | | ● | | | | WP_1078164191<br>WP_2458902941 |
| **Dinoroseobacter_shibae__DFL_12** | ● | | | ● | | | | WP_0121789001<br>WP_0121771061 |
| **Donghicola_eburneus__DSM_29127** | ● | | ● | | | | | WP_0924638921<br>WP_0924636681 |
| **Donghicola_mangrovi__B5_SW_15** | ● | ● | ● | | | | | WP_1771574421<br>WP_1771573501<br>WP_1771573041 |
| **Donghicola_tyrosinivorans__DSM_100212** | ● | ● | ● | | | | | WP_1062622631<br>WP_1062649311<br>WP_1062660041 |
| **Epibacterium_ulvae__U95** | | ● | | | | | | WP_0902165831 |
| **Falsihalocynthiibacter_arcticus__PAMC_20958** | ● | | | ● | | | | WP_0390009601<br>WP_4179351891 |
| **Falsiruegeria_litorea__CECT_7639** | | ● | | | | | | WP_0857967631 |
| **Falsiruegeria_mediterranea__X** | | ● | | | | | | WP_1087858091 |
| **Hasllibacter_halocynthiae__DSM_29318** | ● | | | | | | | WP_2458837471 |
| **Heliomarina_baculiformis__T40_3** | | ● | | | | | | WP_2383697481 |
| **Jannaschia_aquimarina__DSM_28248** | ● | | ● | | | | | WP_0439170331<br>WP_0439206641 |
| **Jannaschia_donghaensis__CECT_7802** | ● | | | | | | | WP_2456242301 |
| **Jannaschia_faecimaris__DSM_100420** | ● | | | ● | | | | WP_0926475321<br>WP_2445045761 |
| **Jannaschia_helgolandensis__DSM_14858** | ● | | ● | | | | | WP_0927644591<br>WP_0927634391 |

| | P1 | P1.1 | P2.1 | P2.2 | U1 | U2 | U3 | Ref |
|---|---|---|---|---|---|---|---|---|
| **Species** | | | | | | | | |
| **Jannaschia_marina__SHC163** | ● | | | | | | | WP_1793806361 |
| **Jannaschia_maritima__LMIT008** | ● | | | | | | | WP_3742135811 |
| **Jannaschia_ovalis__GRR_S6_38** | ● | | ● | | | | | WP_2799658851<br>WP_2799637491 |
| **Jannaschia_pagri__AI_62** | ● | | | | | | | WP_2207471071 |
| **Jannaschia_pohangensis__DSM_19073** | ● | | | | | | | WP_0927823881 |
| **Jannaschia_rubra__DSM_16279** | ● | | | ● | | | | WP_0749626121<br>WP_0556839571 |
| **Jannaschia_seohaensis__DSM_25227** | ● | | ● | | | | ● | WP_1095647431<br>WP_1095638631<br>WP_1095622901 |
| **Leisingera_aquaemixtae__R2C4** | | ● | | | | | | WP_1418906411 |
| **Leisingera_aquimarina__DSM_24565** | | ● | | | | | | WP_0272593951 |
| **Leisingera_caerulea__S122** | | ● | | | | | | WP_2599589411 |
| **Leisingera_daeponensis__DSM_23529** | | ● | | | | | | WP_0085535651 |
| **Leisingera_methylohalidivorans__DSM_14336;_MB2** | | ● | | | | | | WP_0240895721 |
| **Leisingera_thetidis__BMJM1** | | ● | | | | | | WP_2642101511 |
| **Lentibacter_algarum__DSM_24677** | | ● | | | | | | WP_0898882051 |
| **Litoreibacter_albidus__DSM_26922** | ● | | | | | | | WP_0899444941 |
| **Litoreibacter_arenae__DSM_19593** | ● | | | ● | | | | WP_0210999181<br>WP_0211013981<br>WP_0211028081 |
| **Litoreibacter_ascidiaceicola__DSM_100566** | ● | | | ● | | | | WP_0731397121<br>WP_0835886751 |
| **Litoreibacter_halocynthiae__DSM_29467** | ● | | | | | | | WP_1340139331 |
| **Litoreibacter_janthinus__DSM_26921** | ● | | | | | | | WP_0902182331 |
| **Litoreibacter_meonggei__DSM_29466** | ● | | | | | | | WP_1210212371 |
| **Litoreibacter_ponti__DSM_100977** | ● | | | ● | | | | WP_1078462511<br>WP_1078474961 |
| **Litoreibacter_roseus__K6** | ● | | | ● | | | | WP_1598089221<br>WP_2431449171 |
| **Loktanella_agnita__R10SW5** | ● | | | | | | | WP_4116464451 |
| **Loktanella_atrilutea__DSM_29326** | ● | | | | | | | WP_0728572651 |
| **Loktanella_gaetbuli__TSTF_M6** | ● | | | | | | | WP_2267483821 |
| **Lutimaribacter_marinistellae__KCTC_42911** | | ● | ● | | | | | WP_3867367381<br>WP_3867361341 |
| **Lutimaribacter_pacificus__DSM_29620** | | ● | | | | | | WP_1497888111 |
| **Lutimaribacter_saemankumensis__DSM_28010** | | ● | ● | | | | | WP_0900281631<br>WP_0900308121 |
| **Mameliella_alba__JL351** | | ● | ● | | | | | WP_0431382691<br>WP_0746220521 |
| **Mameliella_sediminis__DP3N28_2** | | ● | ● | | | | | WP_2184510251<br>WP_2184512761<br>WP_2184502291 |
| **Marinovum_algicola__FF3** | | ● | | | | | | WP_0748385901 |
| **Maritimibacter_alexandrii__LZ_17** | ● | | | | | | | WP_1384241211 |
| **Maritimibacter_alkaliphilus__HTCC2654** | ● | | | | | | | WP_0083296371 |
| **Maritimibacter_dapengensis__DP4N28_5** | ● | | | | | | | WP_2183910811 |
| **Marivita_cryptomonadis__CL_SK44** | | ● | | | | | | WP_0856292731 |
| **Marivita_geojedonensis__DPG_138** | | ● | | | | | | WP_0856415211 |
| **Marivita_hallyeonensis__DSM_29431** | | ● | | | | | | WP_0727767581 |
| **Mesobacterium_hydrothermale__TK19101** | | ● | | | | | | WP_3262956201 |
| **Mesobaculum_littorinae__M0103** | ● | | | | | | | WP_1279046591 |
| **Nereida_ignava__TBRI4** | ● | | | | | | | WP_3914815451 |
| **Oceanicola_granulosus__HTCC2516** | ● | | | | | | | WP_0072538661 |

| | P1 | P1.1 | P2.1 | P2.2 | U1 | U2 | U3 | Ref |
|---|---|---|---|---|---|---|---|---|
| **Species** | | | | | | | | |
| **Oceaniovalibus_guishaninsula__JLT2003** | ● | | | | | | | WP_0074251571 |
| **Octadecabacter_algicola__D2_3** | ● | | | | | | | WP_2352593221 |
| **Octadecabacter_antarcticus__307** | ● | | | | | | | WP_0154993051 |
| **Octadecabacter_arcticus__238** | ● | | | | | | | WP_0154941451 |
| **Octadecabacter_ascidiaceicola__CECT_8868** | ● | | | | | | | WP_0939959301 |
| **Octadecabacter_dasysiphoniae__G9_8** | ● | | | | | | | WP_2352242311 |
| **Octadecabacter_temperatus__SB1** | ● | | | | | | | WP_0498339481 |
| **Pacificibacter_marinus__CECT_7971** | ● | | | ● | | | | WP_0858493821<br>WP_0858495461 |
| **Pacificibacter_maritimus__DSM_104731** | ● | | | | | | | WP_1237912631 |
| **Pacificoceanicola_onchidii__XY_301** | | ● | | | | | | WP_1364392081 |
| **Palleronia_abyssalis__CECT_8504** | ● | | | | | | | WP_1088951221 |
| **Palleronia_aestuarii__DSM_22009** | ● | | | | | | | WP_1115361111 |
| **Palleronia_caenipelagi__JBTF_M29** | ● | | | | | | | WP_1428332421 |
| **Palleronia_pelagia__DSM_26893** | ● | | | | | | | WP_0918454061 |
| **Palleronia_rufa__MOLA_401** | ● | | | | | | | WP_0361772021 |
| **Palleronia_salina__DSM_26892** | ● | | | | | | | WP_0731282531 |
| **Parasulfitobacter_algicola__1151** | ● | | | | | | | WP_1741358891 |
| **Pelagimonas_phthalicica__DSM_26923** | | ● | | | | | | WP_0992438771 |
| **Pelagimonas_varians__DSM_23678** | | ● | ● | ● | | | | WP_0978066841<br>WP_0978047391<br>WP_0978067471 |
| **Phaeobacter_gallaeciensis__DSM_26640** | | ● | | | | | | WP_0240967671 |
| **Phaeobacter_inhibens__DSM_17395** | | ● | | | | | | WP_0148804101 |
| **Phaeobacter_inhibens__P54** | | ● | | | | | | WP_1028768511 |
| **Phaeobacter_italicus__DP7Y7_1** | | ● | | | | | | WP_2224596251 |
| **Phaeobacter_marinintestinus__UB_M7** | | ● | | ● | | | | WP_1463450311<br>WP_1463469261 |
| **Phaeobacter_piscinae__S26** | | ● | | | | | | WP_0401705421 |
| **Phaeobacter_porticola__P97** | | ● | | | | | | WP_0725047951 |
| **Phycobacter_azelaicus__F10** | | ● | | | | | | WP_1929662111 |
| **Pontibaca_methylaminivorans__DSM_21219** | | ● | | ● | | | | WP_0766494171<br>WP_0839460301 |
| **Pontibaca_salina__S1109L** | | ● | | | | | | WP_1986859911 |
| **Ponticoccus_alexandrii__C31** | | ● | | | | | | WP_0238525501<br>WP_0396157831 |
| **Ponticoccus_litoralis__KCCM_90028** | | ● | | | | | | WP_3471680161 |
| **Poseidonocella_pacifica__DSM_29316** | | ● | | ● | | | | WP_0920634421<br>WP_2176459311 |
| **Poseidonocella_sedimentorum__KMM_9023,NRIC_0796,JCM_17311,KCTC_23692** | | ● | | | | | | WP_0920762961 |
| **Primorskyibacter_aestuariivivens__OITF_36** | | ● | | | | | ● | WP_2714196951<br>WP_2714216611 |
| **Primorskyibacter_flagellatus__CGMCC_1.12664** | | ● | | | | | | WP_1884762721 |
| **Primorskyibacter_marinus__PX7** | | ● | | | | | | WP_1165986571 |
| **Primorskyibacter_sedentarius__DSM_104836** | | ● | | ● | | | | WP_1322413371<br>WP_1322410721<br>WP_1322452361 |
| **Profundibacterium_mesophilum__KAUST100406_0324** | ● | | | | | | | WP_1599642071 |
| **Pseudoponticoccus_marisrubri__SJ5A_1** | | ● | ● | | | | | WP_0588641921<br>WP_2404845001 |
| **Pseudoruegeria_aquimaris__CECT_7680** | ● | | | | | | | WP_1398385291 |
| **Pseudosulfitobacter_koreensis__AP_MA_4** | | ● | | | | | | WP_2582957361 |
| **Pseudosulfitobacter_pseudonitzschiae__H46** | | ● | | | | | | WP_0379262251 |

| | P1 | P1.1 | P2.1 | P2.2 | U1 | U2 | U3 | Ref |
|---|---|---|---|---|---|---|---|---|
| **Species** | | | | | | | | |
| **Roseicyclus_amphidinii__Amp_Y_6** | ● | | | | | | | WP_2842632681 |
| **Roseicyclus_elongatus__DFL_43** | ● | | ● | | | | | WP_0253106071<br>WP_0253108311 |
| **Roseicyclus_marinus__CCMM001** | ● | | | | | | | WP_2778242881 |
| **Roseicyclus_persicicus__KMU_115** | ● | | | | | | | WP_1686218851 |
| **Roseicyclus_sediminis__SDUM158016** | ● | | ● | | | | | WP_2628793351<br>WP_2628773341 |
| **Roseivivax_isoporae__LMG_25204** | | ● | | | | | | WP_0437670921 |
| **Roseivivax_jejudonensis__CECT_8625** | | ● | | | | | | WP_0857931751 |
| **Roseivivax_lentus__DSM_29430** | | ● | ● | | | | | WP_0764456511<br>WP_2349903321 |
| **Roseivivax_marinus__DSM_27511** | | ● | | | | | | WP_0928098451 |
| **Roseobacter_cerasinus__AI77** | | ● | | ● | | | | WP_1599791121<br>WP_2751136831 |
| **Roseobacter_denitrificans__FDAARGOS_309** | | ● | | | | | | WP_0115681161 |
| **Roseobacter_fucihabitans__B14** | | ● | | | | | | WP_1874304831 |
| **Roseobacter_insulae__YSTF_M11** | | ● | | ● | | | | WP_2195066451<br>WP_2194977941 |
| **Roseobacter_litoralis__Och_149** | | ● | | | | | | WP_0139609351 |
| **Roseobacter_ponti__DSM_106830** | | ● | | | | ● | | WP_1696412981<br>WP_1696411441 |
| **Roseobacter_sinensis__WL0113** | | ● | | ● | | | | WP_2638427561<br>WP_2638441901<br>WP_2638438211 |
| **Roseobacter_weihaiensis__H9** | | ● | | ● | | | | WP_2272674201<br>WP_2272693901<br>WP_2272694951 |
| **Roseovarius_aestuarii__KCTC_22174** | | ● | | ● | | | | WP_0858004641<br>WP_0857983161 |
| **Roseovarius_aestuariivivens__GHTF_24** | | ● | | | | | | WP_1355036671 |
| **Roseovarius_albus__CECT_7450** | | ● | | | | | | WP_0858057271 |
| **Roseovarius_aquimarinus__CAU_1059** | | ● | | | | | | WP_3771700671 |
| **Roseovarius_arcticus__MK6_18** | | ● | | ● | | | | WP_1389344901<br>WP_1389344251 |
| **Roseovarius_atlanticus__R12B** | | ● | | | | | | WP_0577901001 |
| **Roseovarius_azorensis__DSM_100674** | | ● | | | | | | WP_0930346781 |
| **Roseovarius_carneus__LXJ103** | | ● | | | | | | WP_1093876061 |
| **Roseovarius_conchicola__2305UL8_3** | | ● | | | | | | WP_3712266451 |
| **Roseovarius_confluentis__SAG6** | | ● | | ● | | | | WP_1037616341<br>WP_1037640791 |
| **Roseovarius_dicentrarchi__YLY04** | | ● | | ● | ● | | | WP_1139122651<br>WP_1139124191<br>WP_1624973451 |
| **Roseovarius_faecimaris__MME_070** | | ● | | | | | | WP_1577064821 |
| **Roseovarius_gaetbuli__CECT_8370** | | ● | ● | ● | | | | WP_0858265841<br>WP_1398380691<br>WP_3063723821 |
| **Roseovarius_halotolerans__DSM_29507** | | ● | | | | | | WP_0858180311 |
| **Roseovarius_indicus__DSM_26383** | | ● | ● | | | | | WP_0578187241<br>WP_0749398881 |
| **Roseovarius_litoreus__DSM_28249** | | ● | | | | | | WP_1497792711 |
| **Roseovarius_litorisediminis__CECT_8287** | | ● | | ● | | | | WP_0858930231<br>WP_2358622251 |
| **Roseovarius_lutimaris__DSM_28463** | | ● | | ● | | | | WP_0928356221<br>WP_2457364301 |
| **Roseovarius_mucosus__DSM_17069** | | ● | ● | | | | | WP_0372721171<br>WP_0372744651 |
| **Roseovarius_nanhaiticus__DSM_29590** | | ● | | | | | | WP_0765335221 |
| **Roseovarius_nubinhibens__ISM** | | ● | | | | | | WP_0098148151 |

| | P1 | P1.1 | P2.1 | P2.2 | U1 | U2 | U3 | Ref |
|---|---|---|---|---|---|---|---|---|
| **Species** | | | | | | | | |
| **Roseovarius_pacificus__DSM_29589** | | ● | ● | | | | | WP_0730327351<br>WP_2297095771 |
| **Roseovarius_pelagicus__HL_MP18** | | ● | | | ● | | | WP_2630486501<br>WP_2630474601 |
| **Roseovarius_phycicola__S88** | | ● | | | | | | WP_3385485781 |
| **Roseovarius_rhodophyticola__W115** | | ● | | ● | | | | WP_3170576521<br>WP_3391067301 |
| **Roseovarius_spongiae__HN_E21** | | ● | | | | | | WP_1211634941 |
| **Ruegeria_alba__1NDH52C** | | ● | | | | | | WP_2389047801 |
| **Ruegeria_aquimaris__XHP0148** | | ● | | ● | | | | WP_2638269991<br>WP_2638300711 |
| **Ruegeria_arenilitoris__HKCCA0515** | | ● | | | | | | WP_1703665051 |
| **Ruegeria_atlantica__CECT_4292** | | ● | | | | | | WP_0582758911 |
| **Ruegeria_conchae__TW15** | | ● | | | | | | WP_0104391461 |
| **Ruegeria_denitrificans__CECT_5091** | | ● | | | | | | WP_0582806991 |
| **Ruegeria_faecimaris__DSM_28009** | | ● | | | | | | WP_1426330001 |
| **Ruegeria_haliotis__B1Z28** | | ● | | | | | | WP_1768634201 |
| **Ruegeria_halocynthiae__DSM_27839** | | ● | | | | | | WP_0747352271 |
| **Ruegeria_intermedia__DSM_29341** | | ● | | ● | | | | WP_1497760981<br>WP_1497741551 |
| **Ruegeria_lacuscaerulensis__HKCCC1251** | | ● | | | | | | WP_1707888871 |
| **Ruegeria_marina__CGMCC_1.9108** | | ● | | ● | | | | WP_0930301441<br>WP_0930267291 |
| **Ruegeria_marisflavi__WL0004** | | ● | | | | | | WP_2633879331 |
| **Ruegeria_marisrubri__ZGT118** | | ● | | | | | | WP_0683434971 |
| **Ruegeria_meonggei__R78036** | | ● | | | | | | WP_3771867231 |
| **Ruegeria_pomeroyi__DSS_3** | | ● | | ● | | | | WP_0110470331<br>WP_0110458851 |
| **Ruegeria_profundi__DP1N0_1** | | ● | ● | | | | | WP_2248893641<br>WP_2248899491 |
| **Ruegeria_sediminis__CAU_1488** | | ● | ● | | | | | WP_1388396391<br>WP_1388399841 |
| **Ruegeria_sp.__TM1040** | | ● | | | | | | WP_0115390431 |
| **Sagittula_marina__DSM_102235** | | ● | | | | | | WP_1839633631 |
| **Sagittula_salina__M10.9X** | | ● | | | | | | WP_2093634631<br>WP_2093588961 |
| **Sagittula_stellata__E_37** | | ● | | | | | | WP_0058566051 |
| **Salibaculum_halophilum__WDS1C4** | ● | | | | | | | WP_0848605741 |
| **Salipiger_abyssi__JLT2014** | | ● | | ● | | | | WP_0766961531<br>WP_0766971801 |
| **Salipiger_aestuarii__AD8** | | ● | | ● | | | | WP_0095052481<br>WP_0344536131 |
| **Salipiger_bermudensis__SY_1_12** | | ● | | ● | | | | WP_2070763681<br>WP_2070762891<br>WP_3406891781 |
| **Salipiger_mangrovisoli__6D45A** | | ● | ● | | | | | WP_1941368131<br>WP_1941361621 |
| **Salipiger_marinus__VSW210** | | ● | | | | | | WP_3247827761 |
| **Salipiger_pallidus__CGMCC_1.15762** | | ● | | | | | | WP_1887902291 |
| **Salipiger_pentaromativorans__P9** | | ● | | ● | | | | WP_2580928111<br>WP_2580931121 |
| **Salipiger_profundus__JLT2016** | | ● | | ● | | | | WP_0766239321<br>WP_0766234841 |
| **Salipiger_thiooxidans__DSM_10146** | | ● | | ● | | | | WP_0403839011<br>WP_0088849261 |
| **Seohaeicola_nanhaiensis__CGMCC_1.12759** | | ● | | ● | | | | WP_3807159251<br>WP_4177650751 |
| **Seohaeicola_saemankumensis__CCUG_55328** | | ● | ● | | | | | WP_3807894141<br>WP_3807946481 |

| | P1 | P1.1 | P2.1 | P2.2 | U1 | U2 | U3 | Ref |
|---|---|---|---|---|---|---|---|---|
| **Species** | | | | | | | | |
| **Seohaeicola_zhoushanensis__KCTC_42650** | | ● | | ● | | | | WP_1896828891<br>WP_2298640781 |
| **Shimia_abyssi__DSM_100673** | | ● | | | | | | WP_1066065051 |
| **Shimia_aestuarii__DSM_15283** | | ● | | | | | | WP_0930947481 |
| **Shimia_biformata__JCM_18818** | | ● | | | | | | WP_2041139531 |
| **Shimia_haliotis__DSM_28453** | | ● | | | | | | WP_0933250421 |
| **Shimia_isoporae__DSM_26433** | | ● | | | | | | WP_1328590711 |
| **Shimia_litoralis__CL_ES2** | | ● | | ● | ● | | | WP_1380155061<br>WP_1380153341<br>WP_1380170981 |
| **Shimia_marina__DSM_26895** | | ● | | ● | | | | WP_0582401531<br>WP_0834990031 |
| **Shimia_ponticola__WX04** | ● | | | | | | | WP_1471255471 |
| **Shimia_sagamensis__DSM_29734** | | ● | | | | | | WP_2834261791 |
| **Shimia_sediminis__ZQ172** | | ● | | | | | | WP_1271153511 |
| **Shimia_thalassica__CECT_7735** | | ● | | ● | | | | WP_0583116321<br>WP_2334882561 |
| **Sulfitobacter_aestuarii__TISTR_2562** | | ● | | | | | | WP_3863722131 |
| **Sulfitobacter_aestuariivivens__TSTF_M16** | | ● | | ● | | | | WP_1910737741<br>WP_1910763941 |
| **Sulfitobacter_albidus__JK7_1** | | ● | | | | | | WP_2127053181 |
| **Sulfitobacter_alexandrii__AM1_D1** | | ● | | | | | | WP_0719710461 |
| **Sulfitobacter_brevis__DSM_11443** | | ● | | | | | | WP_0939239991 |
| **Sulfitobacter_delicatus__DSM_16477** | | ● | | | | | | WP_0937397631 |
| **Sulfitobacter_donghicola__KCTC_12864** | | ● | | | | | | WP_0250580831 |
| **Sulfitobacter_dubius__DSM_16472** | | ● | | | | | | WP_0939267631 |
| **Sulfitobacter_faviae__OXR_9** | | ● | | | | | | WP_3223280751 |
| **Sulfitobacter_geojensis__MM_124** | | ● | | | | | | WP_0250452961 |
| **Sulfitobacter_guttiformis__KCTC_32187** | | ● | | ● | | | | WP_0250614851<br>WP_0379682451 |
| **Sulfitobacter_indolifex__DSM_14862** | | ● | | | | | | WP_0407009701 |
| **Sulfitobacter_litoralis__DSM_17584** | | ● | | | | | | WP_0937332561 |
| **Sulfitobacter_marinus__DSM_23422** | | ● | | ● | | | | WP_0939145021<br>WP_0939169401<br>WP_0939154181 |
| **Sulfitobacter_maritimus__S0837** | | ● | | | | | | WP_1748600441 |
| **Sulfitobacter_mediterraneus__SC7_37** | | ● | | | | | | WP_2031995891 |
| **Sulfitobacter_noctilucae__NB_68** | | ● | | | | | | WP_0250535251 |
| **Sulfitobacter_noctilucicola__NB_77** | | ● | | | | | | WP_0250567811 |
| **Sulfitobacter_pacificus__NBRC_109915** | | ● | | | | | | WP_2843730281 |
| **Sulfitobacter_pontiacus__DSM_10014** | | ● | | | | | | WP_0746356051 |
| **Sulfitobacter_profundi__NBRC_113428** | | ● | | | | | | WP_3862839061 |
| **Sulfitobacter_sabulilitoris__HSMS_29** | | ● | | ● | | | | WP_1386634271<br>WP_1386631651 |
| **Sulfitobacter_sediminis__MJW_29** | | ● | | ● | | | | WP_3678766111<br>WP_3678793431 |
| **Sulfitobacter_undariae__DSM_102234** | | ● | | | | | | WP_1845637851 |
| **Tateyamaria_armeniaca__KMU_156** | | ● | | ● | | | | WP_4075938171<br>WP_4075929591 |
| **Tateyamaria_omphalii__KCTC_12333** | | ● | | ● | | | | WP_1893689571<br>WP_1893711351 |
| **Tateyamaria_pelophila__DSM_17270** | | ● | ● | | | | | WP_2234260111<br>WP_2234287511<br>WP_2234287421 |
| **Thalassobacter_stenotrophicus__LXJ116** | ● | | | ● | | | | WP_0581233141<br>WP_1169040001 |

| | P1 | P1.1 | P2.1 | P2.2 | U1 | U2 | U3 | Ref |
|---|---|---|---|---|---|---|---|---|
| **Species** | | | | | | | | |
| **Thalassobius_vesicularis__CC_AMW_E** | | ● | ● | | | | | WP_1363381161<br>WP_1363393241 |
| **Thalassococcus_arenae__CAU_1522** | | ● | ● | | | | | WP_2177784491<br>WP_3485411021 |
| **Thalassococcus_halodurans__DSM_26915** | | ● | | | | | | WP_1039087721 |
| **Thalassococcus_lentus__KCTC_32084** | | ● | | | | | | WP_2714328081 |
| **Thalassococcus_profundi__WRAS1** | | ● | | | | | | WP_1145123771 |
| **Thalassovita_aquimarina__KMM_8518** | | ● | ● | | | | | WP_2127011631<br>WP_2127018511 |
| **Thalassovita_litoralis__DSM_29506** | | ● | ● | | | | | WP_1424950071<br>WP_1424913401 |
| **Thalassovita_mangrovi__GS_10** | | ● | ● | | | | | WP_1609743841<br>WP_1609733581 |
| **Thalassovita_mediterranea__DSM_16398** | | ● | | | | | | WP_0583200161 |
| **Thetidibacter_halocola__KMU_90** | | ● | ● | | | | | WP_2125358741<br>WP_3463465891 |
| **Thiosulfatihalobacter_marinus__GL_11_2** | | ● | | | | | | WP_1979188451 |
| **Tranquillimonas_alkanivorans__DSM_19547** | ● | | | | | | | WP_0934177041 |
| **Tropicibacter_alexandrii__LMIT003** | | ● | | | | | | WP_1216313081 |
| **Tropicibacter_naphthalenivorans__CECT_7648** | | ● | | | | | | WP_0582461321 |
| **Tropicibacter_oceani__YMD87** | | ● | | ● | | | | WP_2823001181<br>WP_2823022281 |
| **Tropicimonas_marinistellae__SF_16** | ● | | ● | | | | ● | WP_0681099101<br>WP_0780593641<br>WP_1614709581 |
| **Tropicimonas_omnivorans__F158** | ● | | | | | | | WP_3116917351 |
| **Tropicimonas_sediminicola__DSM_29339** | ● | | ● | | | | ● | WP_0892309451<br>WP_1764427841<br>WP_1764429971 |
| **Wenxinia_marina__CGMCC_1.6105** | ● | | | | | | | WP_0183015061 |
| **Wenxinia_saemankumensis__DSM_100565** | ● | | | | | | | WP_1393004891 |
| **[Roseibacterium]_beibuensis__JCM_18015** | ● | | ● | | | | | WP_2595502001<br>WP_2595497791 |
